# A Rationally Designed Antimicrobial Peptide from Structural and Functional Insight of *Clostridium difficile* Translation Initiation Factor 1

**DOI:** 10.1101/2023.06.30.547268

**Authors:** Elvira Alanis, Faith Aguilar, Niaz Banaei, Frank B. Dean, Miguel Alanis, James M. Bullard, Yonghong Zhang

**Author notes:** Address correspondence to: Yonghong Zhang, Department of Chemistry, ESCNE 4.620, The University of Texas Rio Grande Valley, 1201 W. University Drive, Edinburg, TX 78539.

## Abstract

A significant increase of hospital-acquired bacterial infections during the COVID-19 pandemic has become an urgent medical problem. *Clostridioides difficile* is an urgent antibiotic-resistant bacterial pathogen and a leading causative agent of nosocomial infections. The increasing recurrence of *C. difficile* infection and antibiotic resistance in *C. difficile* has led to an unmet need for discovery of new compounds distinctly different from present antimicrobials, while antimicrobial peptides as promising alternatives to conventional antibiotics have attracted growing interest recently. Protein synthesis is an essential metabolic process in all bacteria and a validated antibiotic target. Initiation factor 1 from *C. difficile* (Cd-IF1) is the smallest of the three initiation factors that acts to establish the 30S initiation complex to initiate translation during protein biosynthesis. Here we report the solution NMR structure of Cd-IF1 which adopts a typical β-barrel fold and consists of a five-stranded β-sheet and one short α-helix arranged in the sequential order β1-β2-β3-α1-β4-β5. The interaction of Cd-IF1 with the 30S ribosomal subunit was studied by NMR titration for the construction of a structural model of Cd-IF1 binding with the 30S subunit. The short α-helix in IF1 was found to be critical for IF1 ribosomal binding. A peptide derived from this α-helix was tested and displayed a high ability to inhibit the growth of *C. difficile* and other bacterial strains. These results provide a clue for rational design of new antimicrobials.

## Introduction

The coronavirus disease (COVID-19) pandemic has dramatically examined the global healthcare systems resulting in a <u>significant increase of hospital-acquired infections</u> (HAIs) caused by bacterial pathogens. Among HAIs, *Clostridioides difficile* infection (CDI) is one of the most common healthcare-associated infections and one of the most important global public health threats. *C. difficile* is a Gram-positive and spore-forming anaerobic bacillus that produces toxins to cause infectious diseases such as antibiotic-associated diarrhea, pseudomembranous colitis, toxic megacolon, *etc*. [1]. The CDI treatment has become more challenging owing to the rising emergence of new hypervirulent strains, the increasing CDI incidence/recurrence, and antibiotic resistance [2]. This has created an unmet need for the discovery of new antibiotic candidates with new/various modes of action against bacterial pathogens, especially those that cause nosocomial infections and develop multidrug resistance.

Antimicrobial peptides (AMPs) have been proposed as one of the most promising alternatives to antibiotics for the treatment of bacterial infections [3]. AMPs are small peptides with about 12 ∼ 50 amino acids that display antimicrobial activity through various modes of action, which is different from antibiotics with fixed targets. There are more than 3000 AMPs that have been described in the <u>Antimicrobial Peptide Database</u>. Unlike the antibiotics that adopt resistance relatively fast, AMPs develop almost no or limited resistance. In addition, AMPs usually are lower toxic as they are broken down to individual amino acids or small fragments unlike others that might generate harmful metabolites. There are several examples of AMPs that have been investigated in clinical trials, one of which-surotomycin has been used to treat *C. difficile*-associated diarrhea. Recently, a novel peptide-CM-A was reported as an effective inhibitor against *C. difficile* by inducing cell membrane depolarization and permeability [4].

Protein biosynthesis is a fundamental metabolic process occurring in all bacteria. It is a highly dynamic process including initiation, elongation, termination, and ribosomal recycling. Among the four steps of protein biosynthesis, translation initiation is rate limiting, very cooperative, and highly regulated [5]. In prokaryotes, three initiation factors (IF1, IF2, IF3), the messenger RNA (mRNA), and initiator tRNA (fMet-tRNA) assemble with the 30S ribosomal subunit to form the transient 30S initiation complex (30S IC). IF1 is the smallest initiation factor and functions as an essential regulator in the initiation phase during translation. IF1 binds at the A-site of the 30S subunit thereby preventing the initiator tRNA from binding at that site [6]. Structural studies showed that IF1 is similar to the oligonucleotide/oligosaccharide binding fold (OB fold) [6–9]. However, these IF1s demonstrate structural differences in the C-terminal region. For example, Arg^70^ at the C-terminal end of *E. coli* IF1 (PDB ID 1AH9) was identified as critical for IF1 functionality, but the equivalent residue in *Thermus thermophilus* IF1 (PDB ID 1HR0) made no direct contact with the 30S subunit [6, 7]. These results suggest that IF1s in different bacterial species may adopt a distinct structure and interact differently with the 30S ribosomal subunit. In this study, we reported the solution structure of *C. difficile* IF1 (Cd-IF1) and its interaction studies with the 30S ribosomal subunit. The results allowed us to rationally design a short peptide based on Cd-IF1 structure. The peptide was tested and exhibited an inhibitory activity against the growth of *C. difficile* in the tested media and a broad-spectrum antibacterial potential against other bacterial strains. This IF1-derived peptide is likely a new generation of antibacterial candidates.

## Materials and Methods

Oligonucleotides were ordered from Life Technologies Corporation (Carlsbad, CA). The peptides with the amino acid sequence from the short helical region of Cd-IF1 structure were ordered from APeptide Co. Ltd (Shanghai, China). All other chemicals except as indicated below were obtained from Thermo Fisher Scientific (Waltham, MA). DNA sequencing was performed by Functional Bioscience, Inc. (Madison, WI).

### DNA plasmid construction of Clostridioides difficile IF1

The gene encoding *C. difficile* IF1 protein (Cd-IF1, 72 amino acids) was amplified from *C. difficile* genomic DNA [10]. The polymerase chain reaction (PCR) was conducted on Bio-Rad MJ Mini Thermo Cycler using the forward primer (5’-GGCTAGCATGGCCAAAAAAGATGTTATAG −3’) with an *Nhe*I restriction site and the reverse primer (5’-CTGCTCGAGCTTCTTACGCCAAGTAATTC −3’) with an *Xho*I restriction site. The PCR product was subcloned and inserted into a pET-24b (+) plasmid (Novagen) digested with *Nhe*I/*Xho*I placing the gene upstream of a sequence encoding six histidine residues. The constructed DNA plasmid of pET24b Cd-IF1 contains three extra amino acids (MAS) at the N-terminus and a C-terminal six histidine tag (LEHHHHHH). The recombinant plasmid DNA was verified using DNA sequencing provided by Functional Bioscience, Inc. (Madison, Wisconsin). The pET24b Cd-IF1 plasmid was subsequently transformed into One Shot™ BL21(DE3) Chemically Competent *Escherichia coli* (*E. coli*) (Thermo Fisher Scientific) for expression of the recombinant proteins.

### Preparation of Clostridioides difficile IF1

The recombinant Cd-IF1 proteins were over-expressed using the *E. coli* expression system with the induction of isopropyl β-D-1-thiogalactopyranoside (IPTG). Unlabeled proteins were over-expressed in LB media and purified following the standard His-tag protein purification protocol with an additional purification of size-exclusion chromatography. To produce IF1 for use in NMR structural studies, uniformly ^15^N-labeled and ^13^C/^15^N-labeled Cd-IF1 were over-expressed in M9 media with ^15^NH_4_Cl and ^15^NH_4_Cl/^13^C-glucose (Cambridge Isotope Laboratories, Inc. Andover, MA) using the high-yield protein expression protocol [11] and purified as previously described [12]. The proteins were purified to > 98% homogeneity with a molecular weight of 9.5 kDa confirmed by SDS-PAGE. The final proteins were concentrated and exchanged to the potassium phosphate buffer (20 mM KH_2_PO_4_ (pH 5.1), 100 mM KCl, 2.5 mM DTT) using Amicon Ultra-15 Centrifugal Filters (Millipore #UFC900324, 3 kDa cut-off).

### NMR spectroscopy

Cd-IF1 proteins (unlabeled, ^15^N-or ^15^N/^13^C-labeled) were exchanged into phosphate buffer (20 mM potassium phosphate (pH 5.1), 100 mM KCl, 2.5 mM DTT-*d*_10_) with either 8% or 100% D_2_O. A D_2_O-exchanged sample was made for H-D exchange experiments by freezing Cd-IF1 proteins followed by lyophilization and resuspension in 99.9% D_2_O. All NMR experiments were performed at 298 K on a Bruker AVANCE III Ultrashield Plus 600 MHz spectrometer equipped with a double resonance broad band probe (BBO), or a Bruker AVANCE 700 MHz spectrometer equipped with four independent RF channels and triple resonance cryogenic probe (TCI) with Z-axis pulsed field gradient, deuterium decoupling capability, a variable temperature controller. The NMR chemical shift assignments were completed by analyzing the spectra including HNCACB/CBCA(CO)NH, HNCO, HBHA(CO)NH, and ^15^N-HSQC-TOCSY (mixing time of 60 ms). The side chain aliphatic ^1^H and ^13^C resonances were assigned according to ^13^C-CT-HSQC, ^13^C-HCCH-TOCSY, and CCH-TOCSY spectra. For the assignments of aromatic side chains, ^13^C-CT-HSQC-TOCSY, ^13^C-HSQC-NOESY (mixing time of 120 ms) spectra along with 2D ^1^H-^1^H NOESY (mixing time of 100 ms) and TOCSY (mixing time of 100 ms), were used. Stereospecific assignments of chiral methyl groups of valine and leucine were obtained by analyzing ^1^H-^13^C HSQC experiments performed on a protein sample containing 10% ^13^C-labeled Cd-IF1 [13]. The NMR data were processed using NMRPipe [14] and analyzed using Sparky [15].

### NMR structural calculation

Using the NMR resonance assignments of Cd-IF1 [12], three NOESY spectra including ^15^N-edited and ^13^C-edited 3D NOESY-HSQC as well as 2D ^1^H-^1^H NOESY were analyzed to determine NOE-based interproton distances throughout the protein. Backbone torsion angles (*φ* and ψ) were predicted by TALOS-N according to the NMR chemical shift assignments. Hydrogen bonds were verified by identifying slowly exchanging amide protons in hydrogen-deuterium exchange experiments. The NMR-derived distances plus dihedral angles and hydrogen bonds then served as constraints for calculating the three-dimensional structure using distance geometry and restrained molecular dynamics. Protein structure calculations were performed using Xplor-NIH 2.38 following a simulated annealing protocol [16], as previously described [9]. A total of 835 interproton distance relationships, 33 hydrogen bond distances, and 112 dihedral angles (see Table I) were used as restraints included in the structure calculation of Cd-IF1. Fifty independent structures were calculated, and after refinement, the energy-lowest 15 structures were selected and analyzed. The average total and experimental distance energies were 1481.1 ± 6.7 and 100.4 kcal•mol^-1^. The average root-mean-square (rms) deviation from an idealized geometry for bonds and angles were 0.0082 Å and 1.94°. The NMR-derived structures of Cd-IF1 were assessed by PRO-CHECK. The final NMR ensemble of 15 structures with the lowest energy has been deposited in the <u>RCSB Protein Data Bank</u> (**PDB ID 6C00**).

**Table I:** Structure Calculation Statistics of NMR-derived Structures of Cd-IF1

|  |  |
| --- | --- |
| NOE restraints (total) | 835 |
| intra ($i - j = 0$) | 342 |
| medium ($1 \leq i - j \leq 4$) | 298 |
| long ($i - j > 4$) | 195 |
| dihedral angle restraints ($\varphi$ and $\psi$) | 112 |
| hydrogen bond restraints in β-strand regions | 33 |
| RMSD from ideal geometry |  |
| bond length ($\text{\AA}$) | $0.0082 \pm 0.00015$ |
| bond angles (deg) | $1.94 \pm 0.0045$ |
| Ramachandran plot |  |
| most favored region (%) | 85 |
| allowed region (%) | 13 |
| disallowed region (%) | 2 |
| RMSD from average structure |  |
| β-barrel regions (main chain) ($\text{\AA}$) | $0.28 \pm 0.071$ |
| β-barrel regions (non-hydrogen) ($\text{\AA}$) | $0.94 \pm 0.09$ |

### NMR titration

^15^N-labeled Cd-IF1 pure proteins were exchanged to 20 mM MES buffer (pH 6.0) with 50 mM NaCl, 1 mM EDTA, DTT, and 8% D_2_O using a Millipore Amicon Ultra-15 Centrifugal Filter *Ultracel-3K* (Millipore #UFC900324, 3 kDa cutoff), concentrated to a concentration of 50 μM and transferred to an NMR tube after removal of any precipitates by centrifugation. For NMR titration experiments, the sample of ^15^N-labeled Cd-IF1 was first used to record a 2D ^1^H-^15^N HSQC spectrum with 256 (F1) × 1024 (F2) complex points on a Bruker AVANCE III Ultrashield Plus 600 MHz spectrometer at 298 K. And then the sample was titrated by adding a series of increasing amount of 30S ribosomal subunits (34.6 μM) purified from *Pseudomonas aeruginosa* as reported previously [9]. A 2D HSQC spectrum was recorded in each titration. The total four-titration points in one set of experiments were performed with a final molar ratio of the 30S subunit to Cd-IF1 at 1.0 × 10^-2^. The titration data were processed and analyzed using NMRPipe [14].

### Complex model of 30S subunit with Cd-IF1

The complex model of Cd-IF1 bound with the 30S ribosomal subunit was built according to the NMR titration results and the crystal structure of *T. thermophilus* IF1 in complex with the 30S subunit [6]. Cd-IF1 NMR structure (PDB 6C00) was applied to replace *T. thermophilus* IF1 in the complex structure (PDB 1HR0) by PyMOL (Version 2.4.0a0 Open-Source) [9, 17].

### Cd-IF1 derived peptide

The peptide with amino acid sequence derived from the short α-helix (NH_2_-HISGKLRMNFIRILEGDK-COOH) of *C. difficile* IF1 was ordered from APeptide Co. Ltd. (Shanghai, China). The peptide was chemically synthesized and purified by a preparative HPLC method. HPLC purification was performed using Symmetrix ODS-R (5 μm, 250 × 4.60 mm) column. The peptide was eluted using a gradient of buffer A (0.1% Trifluoroacetic Acid in 100% acetonitrile) and B (0.1% Trifluoroacetic Acid in 100% Water) with a flow rate of 1 mL/min and detected at 220 nm as a single peak via HPLC with >95% purity. The molecular weight of the peptide was confirmed by ESI-MS (Agilent-6125B) as the expected value (2127.5).

### Antimicrobial Activity and MIC Assays of Cd-IF1 Peptide

The antimicrobial activity of Cd-IF1-derived peptide was evaluated using Thermo Fisher Scientific 96-Well Microtiter^TM^ Microplates. The representative bacteria for the tests included Gram-positive strains-*C. difficile* (ATCC 43593), *Staphylococcus epidermidis* (ATCC 12228), *Mycobacterium smegmatis* (ATCC 14468), and *Bacillus cereus* (ATCC 14579), and Gram-negative strains-*P. aeruginosa* (ATCC 47085), *Escherichia coli*, and *Proteus vulgaris*. The *E. coli* strain was BL21(DE3) from Invitrogen (One Shot^TM^ BL21(DE3), Cat. NO. C600003) which is descended from the *E. coli* B strain and commonly used for high-level expression of recombinant proteins. *P. vulgaris* was obtained from The Microbiology Laboratory at Department of Biology, The University of Texas Rio Grande Valley, which have been used in General Microbiology Lab Class. For the preparation of *C. difficile* cultures, a small amount of the strain was streaked on the pre-made Brucella Blood Agar plates with 5% Sheep Blood, Hemin and Vitamin K (Thermo Scientific TM) and then incubated in an anaerobic jar (Sigma-Aldrich Inc.) overnight at 37°C. The bacteria were inoculated from the agar plates to Brain Heart Infusion (BHI) broth (Remel Inc. Lenexa, Kansas) and grew in the anaerobic jar until the OD_600_ value reached the desired reading. For *S. epidermidis* and *B. cereus*, the strains were initially incubated on nutrient agar plates and then grown in nutrient broth media. All other bacteria were grown in Luria-Bertani (LB) agar and broth. The Microtiter plates were set up by transferring 50 μl bacterial cultures with an initial OD_600_ value between 0.08 ∼ 0.13 into each well which contained different concentrations of the peptides, or the control substances (DMSO, ampicillin, kanamycin, metronidazole). The peptides were diluted by serial twofold dilutions from the first well across the plate. DMSO and antibiotics (ampicillin, kanamycin, or metronidazole) were used as the negative and positive control, respectively. The Microtiter plates were incubated at 37 °C overnight and then for examination. The assays for each bacterial strain were performed in triplicate, each MIC represented as the average of three independent results.

### Cytotoxicity assay of Cd-IF1 peptide

The toxicity of Cd-IF1 peptide was examined against human embryonic kidney cells (HEK-293). HEK-293 cell cultures in Dulbecco’s modified Eagle’s media (DMEM) with 10% fetal bovine serum (FBS) and Penicillin-Streptomycin Solution were plated in a 96-well plate with about 20,000 cells per well. The plate was then incubated at 37 °C in the incubator supplied with 5% CO_2_ overnight. The peptides were dissolved in DMSO and diluted to yield a final concentration from 3 mg/ml to 1 μg/ml in the cell cultures for the assay. Dichlorodiphenyltrichloroethane (DDT, a potent inhibitor of human cell culture growth as a comparator) was used as the positive control and DMSO was as negative control. HEK-193 cells in the 96-well plate were treated with the peptide, DDT, or DMSO alone for 18 h prior to the cell proliferation assay. The Trevigen TACS MTT kit (Gaithersburg, MD) was utilized to assess impacts on human cell proliferation and/or viability. A 10 μL of MTT reagent wase added into each well and the plate were then incubated under 5% CO_2_ at 37 °C for another 4 h. Finally, 100 µL of detergent reagent was loaded into each well and incubation was continued for overnight. The plate was examined by BioTek Synergy Multi-Mode Microplate Reader. Samples were conducted in double.

## Results

### NMR-derived structure of Cd-IF1

The ^1^H-^15^N heteronuclear single quantum correlation (HSQC) NMR spectrum of Cd-IF1 shows highly dispersed cross peaks with uniform intensities and narrow peak shapes. The HSQC spectral characteristics indicate that the protein is well-folded and adopts a stable three-dimensional conformation. ^15^N-NMR relaxation analysis (*R*_1_ and *R*_2_) of Cd-IF1 indicates an average rotational correlation time of ∼6 ns and molecular weight of ∼10 kDa, suggesting that Cd-IF1 forms a monomer under NMR solution conditions. NMR chemical shift assignments for Cd-IF1 were reported previously (**BMRB no. 27349**) [12]. These assignments served as a basis for collecting nuclear Overhauser effect (NOE) distances, hydrogen-bonds, and dihedral angle restraints from the NMR experimental data which were used for the atomic-resolution structure calculation as described in the Methods section. Structure calculation and statistical data for the final 15 lowest energy conformers (**Protein Databank accession No. 6C00**) are summarized in **Table I**. The calculated structures were validated by PROCHECK, showing 98% of the residues belong to the most favorable/allowed region in the Ramachandran plot.

The final structures of Cd-IF1 experimentally determined by solution NMR spectroscopy are summarized in **Table I**. As shown in **Figure 1**, the final 15 lowest-energy conformers when superimposed have an overall main chain RMSD (root-mean-square deviation) of 0.28 Å. The residues at both N-terminal and C-terminal ends exhibit random-coil chemical shifts suggesting these regions are disordered and flexible as supported by the heteronuclear ^15^N-NOE experimental results [12]. The solution structure of Cd-IF1 as shown in Figure 1-energy-minimized average structure, consists of five stranded β-sheets and one α-helix: β1 (residues 7-18), β2 (residues 21-26), β3 (residues 29-36), α1 (residues 38-43), β4 (residues 50-58), β5 (residues 64-69). Overall, five strands are arranged in the sequential order of β1-β2-β3-α1-β4-β5 as a β-barrel structure and oriented anti-parallel, except for strands β3 and β5 which are close to each other and parallel.

**Figure 1.**
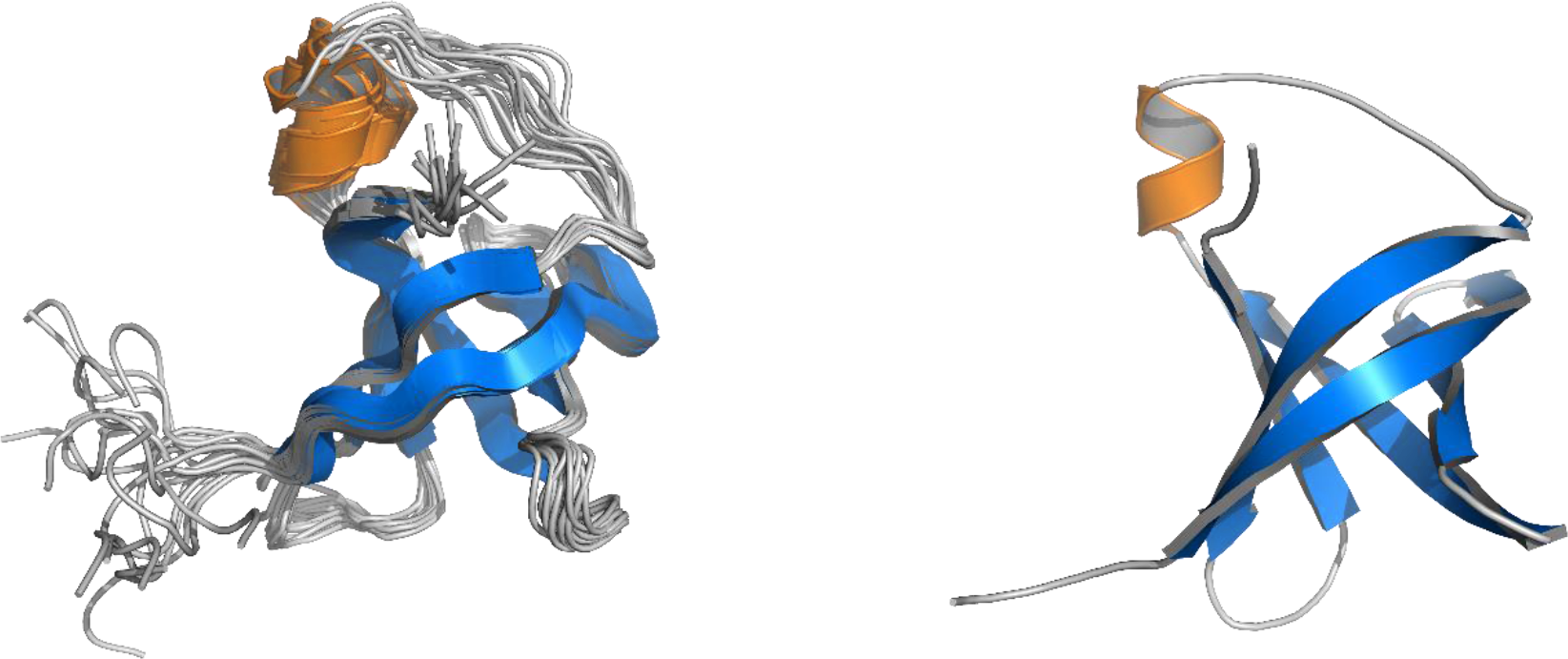
NMR-derived structures of *C. difficile* IF1 in solution. (A) Superposition of main chain atoms of 15 lowest energy structures with RMSD of 0.3 Å (main chain atoms). (B) Ribbon representation of the energy-minimized average main chain structure. The short α-helix is highlighted orange (residues 38-43) and five β-strands blue (β1: residues 7-18, β2: residues 21-26, β3: residues 29-36, β4: residues 50-58, β5: residues 64-69).

While most of the connections between two adjacent β-strands are defined well as typical β-turns (**Figure 1)**, the region between strands β3 and β4 (residues 36-49) contains a short α-helix and appears to be considerably flexible as derived from medium-range NOEs supported by ^15^N-^1^H heteronuclear NOEs [12]. One side of the compact β-barrel is covered by a long flexible loop involving the short helix. The Cd-IF1 adopts the expected OB fold indicative of its binding to the 30S ribosomal subunit [18].

### Surface properties of Cd-IF1

The electrostatic surface representation of Cd-IF1 is shown in **Figure 2**. A positively charged surface composed of strands β3 and β5 close to the short α-helix is rich in basic residues (Arg and Lys), including K39, R41, R46, R64, and R66; while the negatively charged posterior contains mostly acidic residues (Glu and Asp), such as E8, E10, E15, E27, E31, E49, and D51. It is likely that this positively charged surface (**Figure 2)** may make contact with the 30S ribosomal subunit upon binding. Indeed, these residues are conserved (K39, R41, R66) or highly similar (R64) in other bacterial IF1 proteins and are involved in the binding with the 30S ribosomal subunit in the crystal structure of IF1 from *Thermus thermophilus* [6].

**Figure 2.**
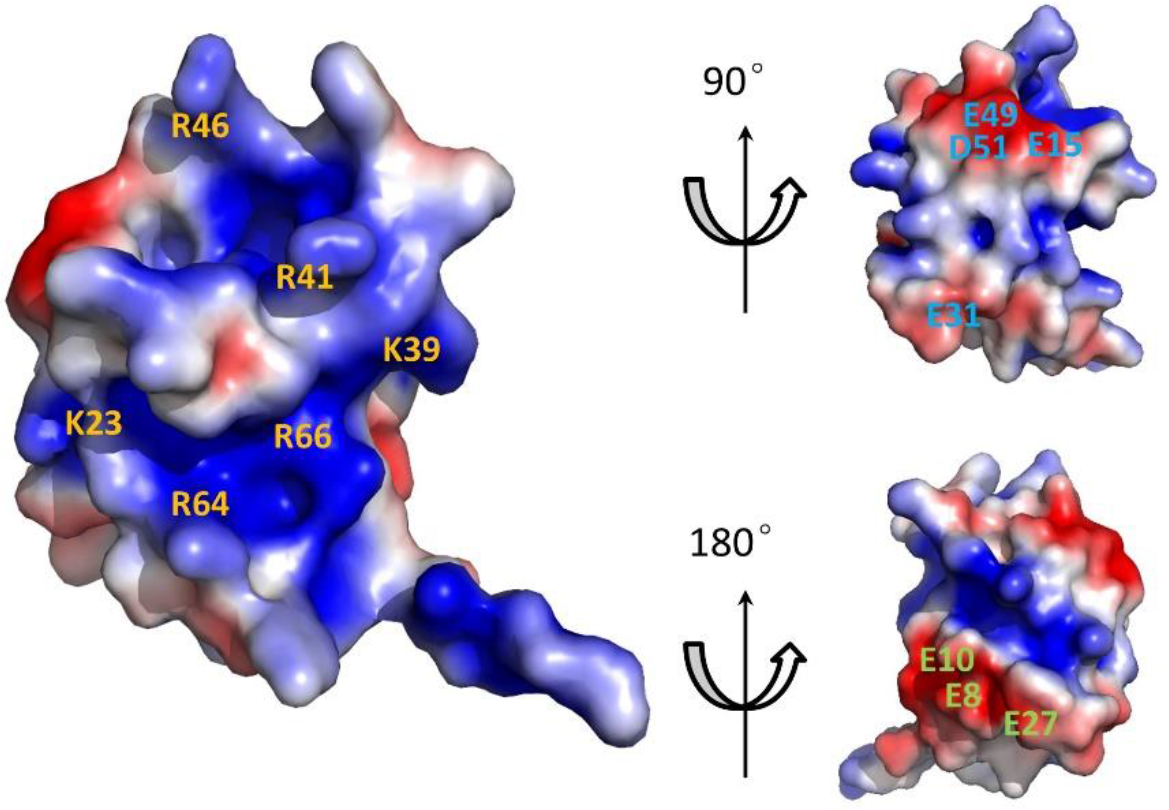
Electrostatic surface potential of *C. difficile* IF1 NMR Structure. The electrostatic potential was calculated using the Adaptive Poisson-Boltzmann Equation (APBS) with ionic strength 0.1 M [30]. The resulting electrostatic potential was visualized by the APBS plugin in PyMOL to generate the surface presentations (±1 kT/e) and views at 90° and 180° rotation around the longitudinal axis. The key charged amino acids are labeled. Red and blue colored regions denote negative and positive charges, respectively.

### Cd-IF1 interaction with the 30S ribosomal subunit

The NMR titration was conducted to examine the binding of Cd-IF1 with the 30S ribosomal subunits. ^15^N-labeled Cd-IF1 proteins were expressed and purified as described in the experimental section. The 30S subunits of *Pseudomonas aeruginosa* ribosomes were prepared according to the procedure reported previously [19], and used for the titration due to the lack of *C. difficile* ribosomes. The ^1^H-^15^N heteronuclear single quantum correlation (HSQC) NMR spectrum of Cd-IF1 in free form exhibited highly dispersed backbone amide cross peaks, uniform peak intensities, and narrow peak shapes, which were typical characteristics of a well-folded protein. The addition of purified 30S subunits induced changes in both chemical shifts and peak intensity in the HSQC spectrum (Figure 3). According to the previous NMR assignments (BMRB no. 27349) [12], the amino acids perturbed by the 30S ribosomes were identified (Figure 3) including M21, H30, H35, I36, K39, V53, V55, G65, R66, W69, etc. These residues with significant perturbations were located on one side of Cd-IF1 β-barrel which was close to the short α-helix.

**Figure 3.**
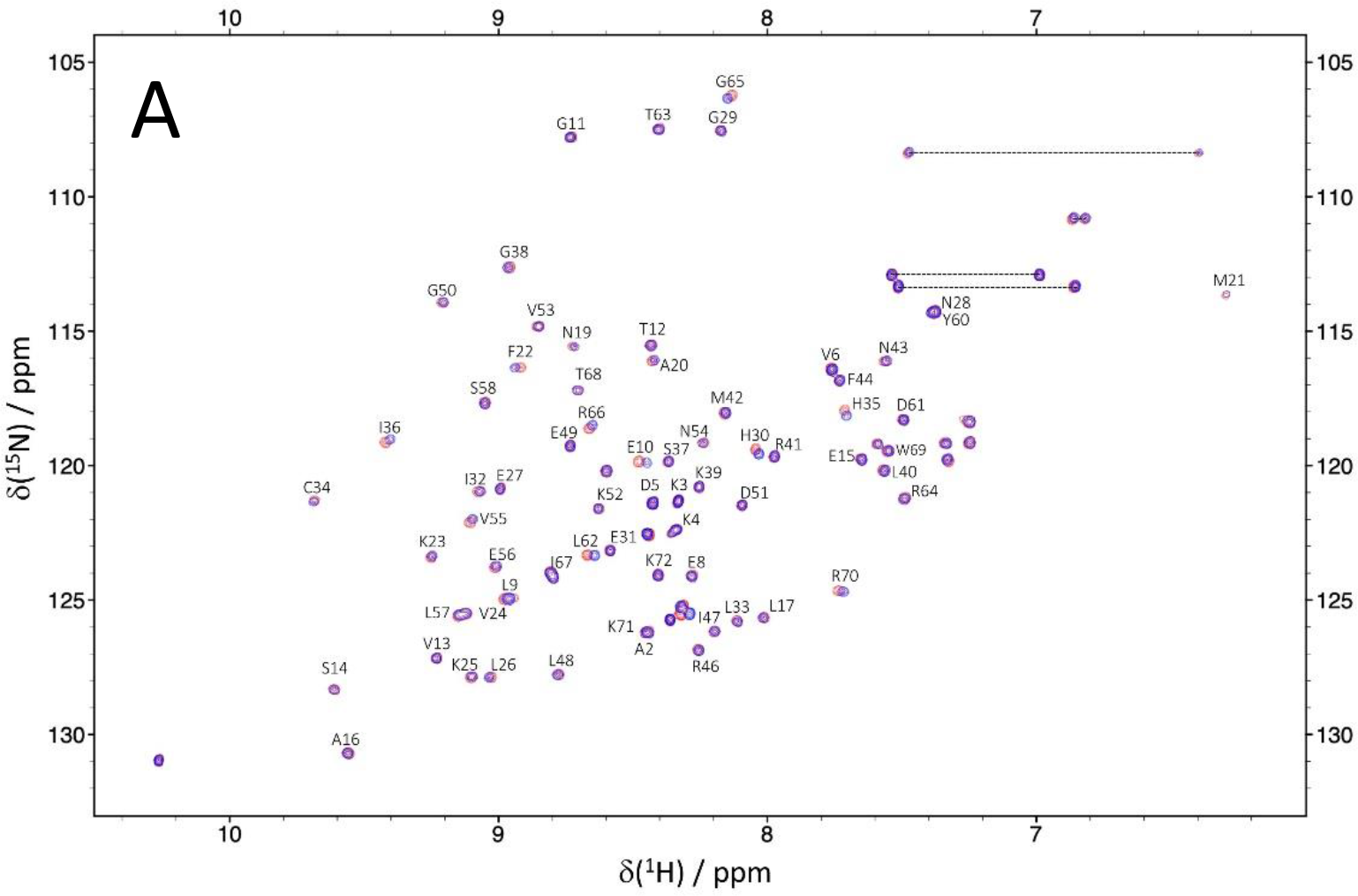
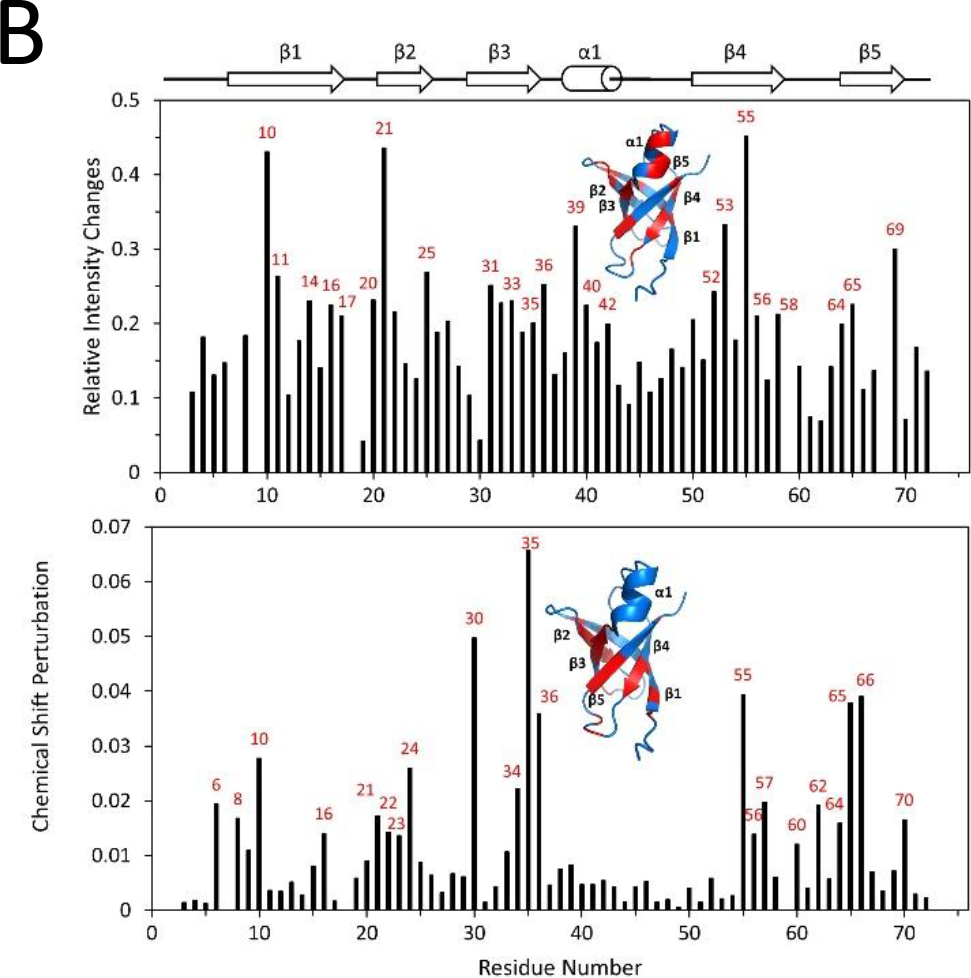
Analysis of ^1^H-^15^N HSQC titrations of ^15^N-labeled Cd-IF1 in 20 mM MES (pH 6.0) with 50 mM NaCl, 1 mM EDTA, DTT, and 8% D2O by the 30S ribosomal subunits. ***A***, Two-dimensional HSQC spectra of Cd-IF1 (50 μM) in the absence (blue) and presence (red) of the 30S with a molar ratio to Cd-IF1 of 1.0 × 10^-2^. The data was collected at 298 K on a Bruker 600 MHz NMR spectrometer. The assigned amino acids are labeled according to the assignments (BMRB accession no. 27349) [12]. ***B***, Histograms of the relative intensity changes (Δ*I*/*I*, up) and the chemical shift perturbation (Δ*δ*, down) of HSQC peaks induced by the 30S, Δ*δ* values were calculated using Equation 1 [31]. 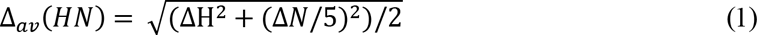 The amino acids exhibiting substantial changes are labeled in the histograms and highlighted in the ribbon diagrams of Cd-IF1 structure (insets).

The NMR titration results were used for computational modeling of the complex structure of Cd-IF1-bound 30S ribosomal subunit according to the crystal structure of *T. thermophilus* IF1 and the 30S (PDB 1HR0) (Figure 4A). Cd-IF1 binds at the A-site of the 30S and makes direct contact with Loop 530, Helix 44, as well as the ribosomal protein S12. The short α-helix in the loop connecting strands three and four at one end of the β-barrel toward the 30S head, is embedded in the groove formed by Loop 530 and Helix 44, with closer to the former. Residues-K39 and M42 tightly interact with the phosphate backbone of Loop 530 (nucleotide G530) (Figure 4B), and residues-R41 and R46 make direct interactions with flipped-out bases, A1492 and A1493. The loop connecting strands one and two inserts into the minor groove of helix 44, and forms contacts with the backbone of several nucleotides. The surface of one side of the β-barrel containing strands three and five faces down to attach the ribosomal protein S12, in which H35, Y60, R64, and R66 were perturbed in the NMR titration.

**Figure 4.**
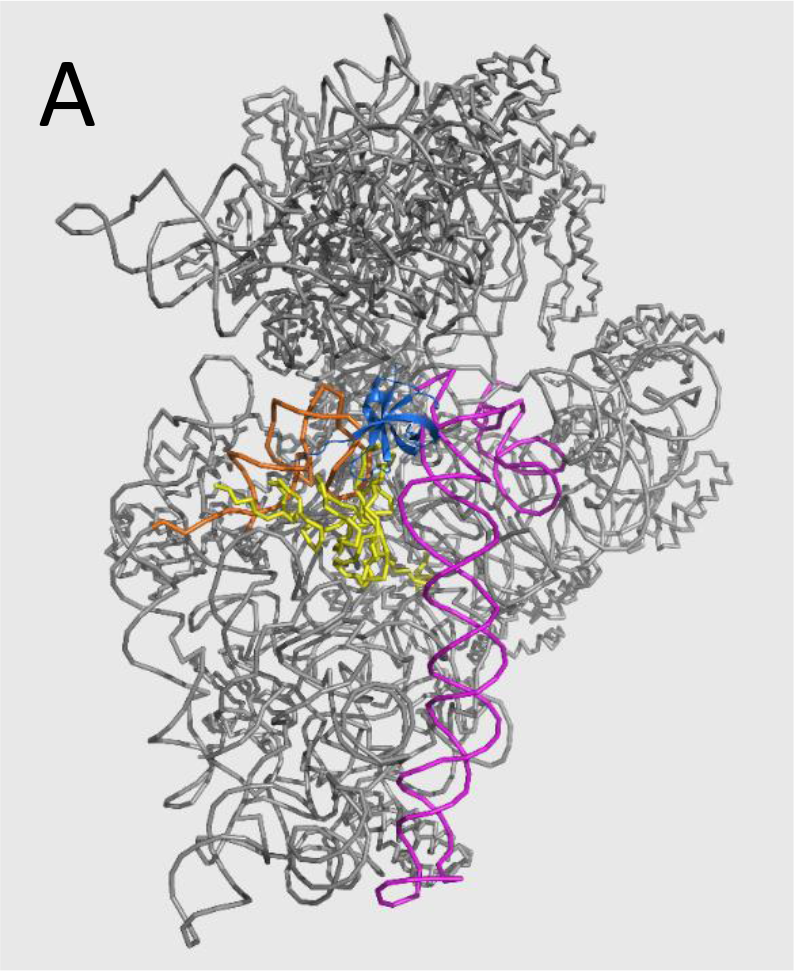
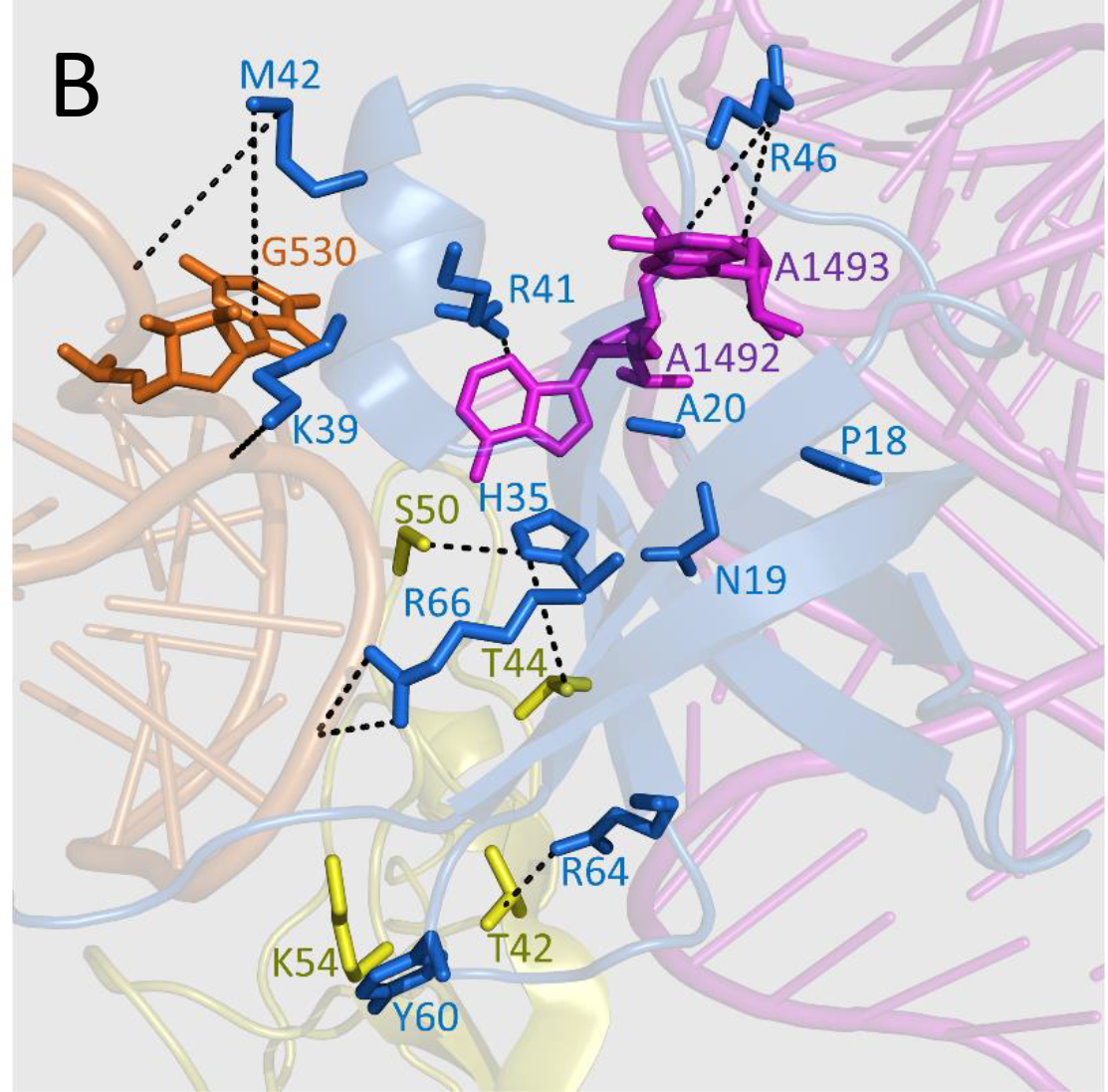
Structure model of Cd-IF1 bound to the 30S ribosomal subunit (PDB 1HR0) constructed according to the NMR titration results. (A) Ribbon representation of the 30S with Cd-IF1 (blue) bound at the A (aminoacyl) site contacting the 530 loop (orange), ribosomal protein S12 (yellow) and helix 44 (magenta). (B) Close-up of Cd-IF1 (blue) bound at the A site of the 30S in ribbon diagram. The key amino acids (blue) of Cd-IF1 with direct contact to the 30S are represented by a stick model with the residue names and numbers labeled. The residues from the 30S interacting with the key amino acids of Cd-IF1 are shown in stick presentation and highlighted in magenta (helix 44), orange (530 loop), and yellow (protein S12).

### Cd-IF1 peptide inhibits C. difficile growth

A short peptide with the amino acid sequence derived from the loop containing the short α-helix was rationally designed to mimic IF1 binding with the 30S subunit for initiation regulation. The peptide was chemically synthesized and used to examine its inhibitory activity against the growth of *C. difficile* in culture media by conducting the Minimum Inhibitory Concentration (MIC) assays.

The MIC determined the lowest concentration of a molecular inhibitor that prevented visible growth of tested microorganisms. An inoculum of *C. difficile* (ATCC 43593) was tested with Cd-IF1-derived peptide at various concentrations using 96-well microtiter plates. The peptides were diluted two-fold to obtain concentrations of 4.0, 2.0, 1.0, 0.5, 0.25, 0.125, 0.063, 0.032, 0.016, 0.008, and 0.004 mg/ml (2.1 μM – 4.0 mM). *C. difficile* inoculum was transferred into BHI broth and cultivated at 37°C under anaerobic conditions, and then the cultures were transferred into a 96-well plate. Metronidazole was used as a positive control. As shown in Figure 5 (Panel A), the peptide showed antimicrobial activity against *C. difficile* at MIC of 0.13 mg/ml (66 μM). Despite being less effective compared with Metronidazole, the peptide exhibited higher inhibitory activity than that of Ampicillin. The latter displayed no inhibition against *C. difficile* under the test conditions (Figure 5A).

**Figure 5.**
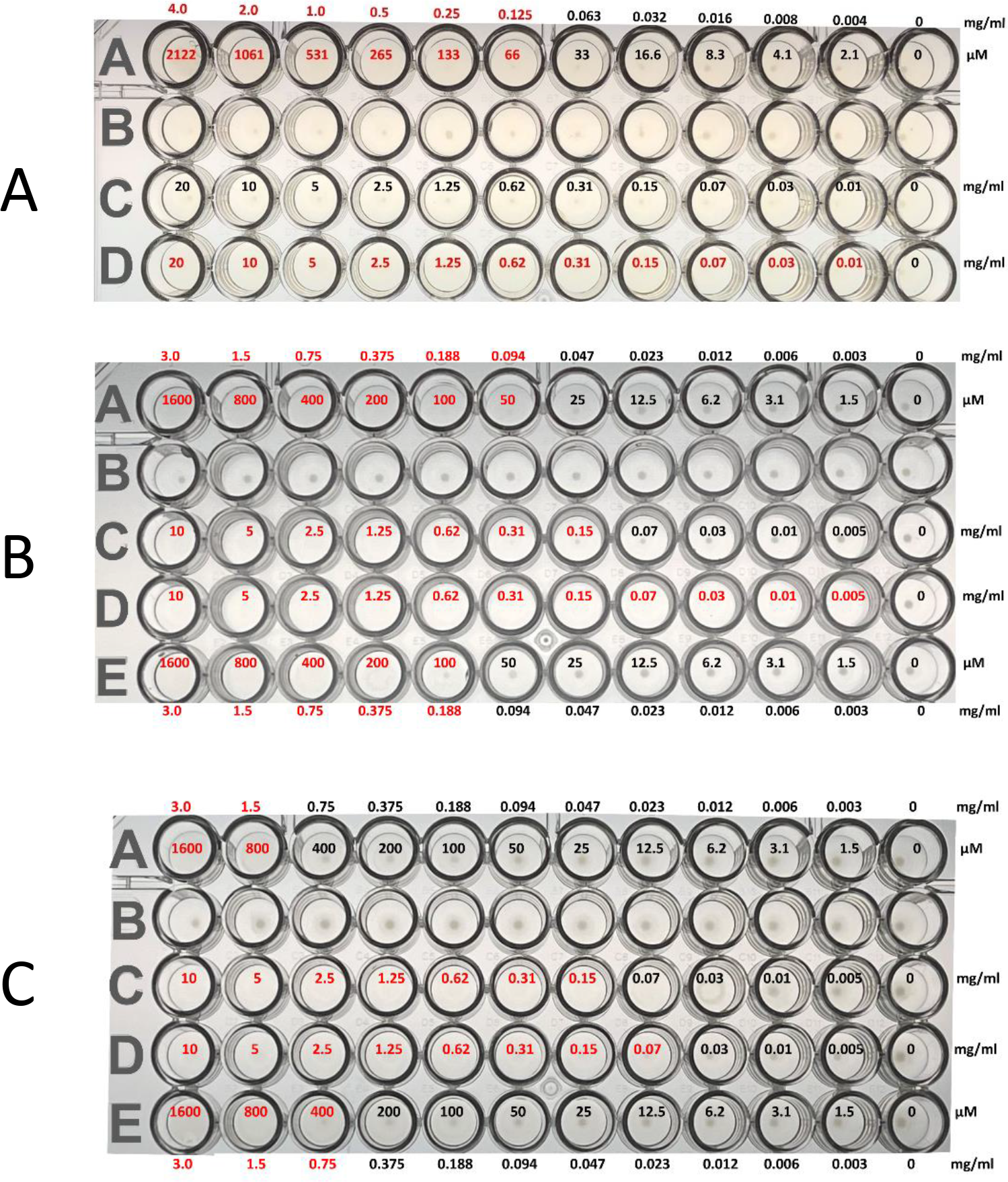
**Panel A**: Cd-IF1 peptide inhibits the growth of *Clostridioides difficile* (ATCC 43593) (row A) in BHI broth with DMSO (row B), ampicillin (row C) and metronidazole (row D) as controls. **Panel B & C**: The peptide shows the inhibition against *Staphylococcus epidermidis* (ATCC 12228) (Panel B - rows A and E) and *Escherichia coli* (BL21(DE3)) (Panel C - rows A and E) in LB media with DMSO (row B in both panels), ampicillin (row C in both panels), and kanamycin (row D in both panel) as controls. The concentration in each well is labeled.

### Cd-IF1 peptide shows broad-spectrum antimicrobial activity but no cytotoxicity

Cd-IF1-derived peptide was further tested in the 96-well MIC assays against other bacteria including Gram-positive and Gram-negative strains. The MIC results of the peptide were summarized in Table II. The peptide inhibited the growth of Gram-positive bacteria-*Staphylococcus epidermidis* (ATCC 12228) with a MIC of 0.14 mg/ml (Figure 5B) and *Mycobacterium smegmatis* (ATCC 14468) with a MIC of 0.37 mg/ml, which were the same or close to the MIC of *C. difficile* under the tested conditions. The peptide also inhibited the growth of *Bacillus cereus* (ATCC 14579) despite a higher MIC value (1.50 mg/ml) compared with that of *C. difficile* (0.13 mg/ml). In addition, the peptide was tested and displayed inhibitory activities against Gram-negative strains, *Escherichia coli* (BL21(DE3)) (Figure 5C), *Proteus vulgaris*, and *Pseudomonas aeruginosa* (ATCC 47085). The values of MICs ranged from 1.0 – 1.5 mg/ml for these Gram-negative species (Table II). These results indicate that the peptide is more active against Gram-positive than Gram-negative bacteria under the tested conditions.

**Table II:**
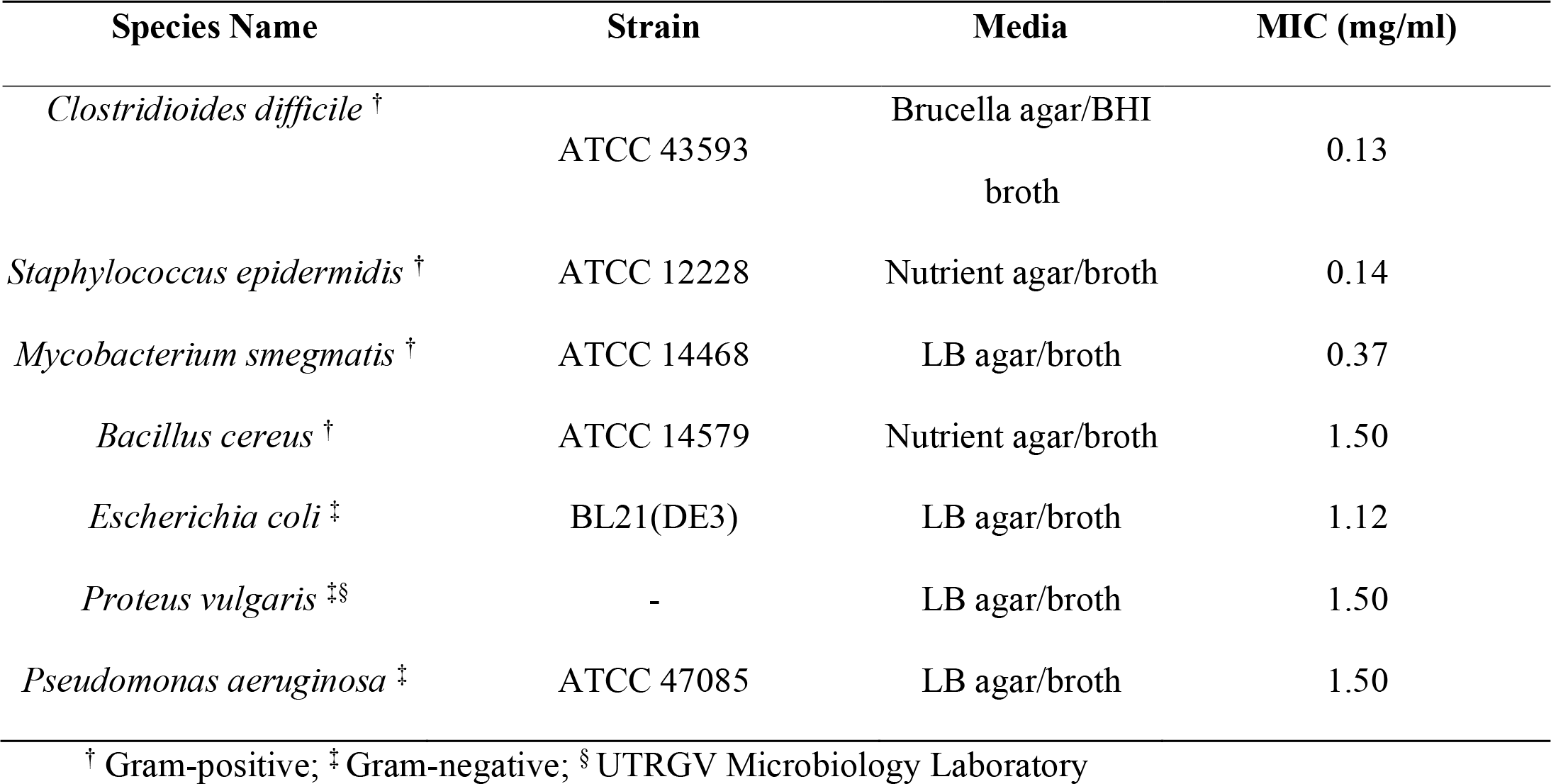
The Minimum Inhibitory Concentrations (MICs) of Cd-IF1 Peptide Against Bacterial Strains

| Species Name | Strain | Media | MIC (mg/ml) |
| --- | --- | --- | --- |
| *Clostridioides difficile* <sup>†</sup> | ATCC 43593 | Brucella agar/BHI<br>broth | 0.13 |
| *Staphylococcus epidermidis* <sup>†</sup> | ATCC 12228 | Nutrient agar/broth | 0.14 |
| *Mycobacterium smegmatis* <sup>†</sup> | ATCC 14468 | LB agar/broth | 0.37 |
| *Bacillus cereus* <sup>†</sup> | ATCC 14579 | Nutrient agar/broth | 1.50 |
| *Escherichia coli* <sup>‡</sup> | BL21(DE3) | LB agar/broth | 1.12 |
| *Proteus vulgaris* <sup>‡§</sup> | - | LB agar/broth | 1.50 |
| *Pseudomonas aeruginosa* <sup>‡</sup> | ATCC 47085 | LB agar/broth | 1.50 |

<sup>†</sup> Gram-positive; <sup>‡</sup> Gram-negative; <sup>§</sup> UTRGV Microbiology Laboratory

In vitro cytotoxicity testing was conducted to evaluate the potentially toxic effect of Cd-IF1-derived peptide on mammalian cells, given that ribosomes are contained within the eukaryotic cells. HEK-293 cells were selected for the test using MTT assays. The cells were treated with 1 – 3000 μg/mL of the peptides for 18 hours under standard tissue culture conditions in duplicate. The peptide was not observed to be toxic to HEK-293 cells at any concentration tested as shown in Figure 6.

**Figure 6.**
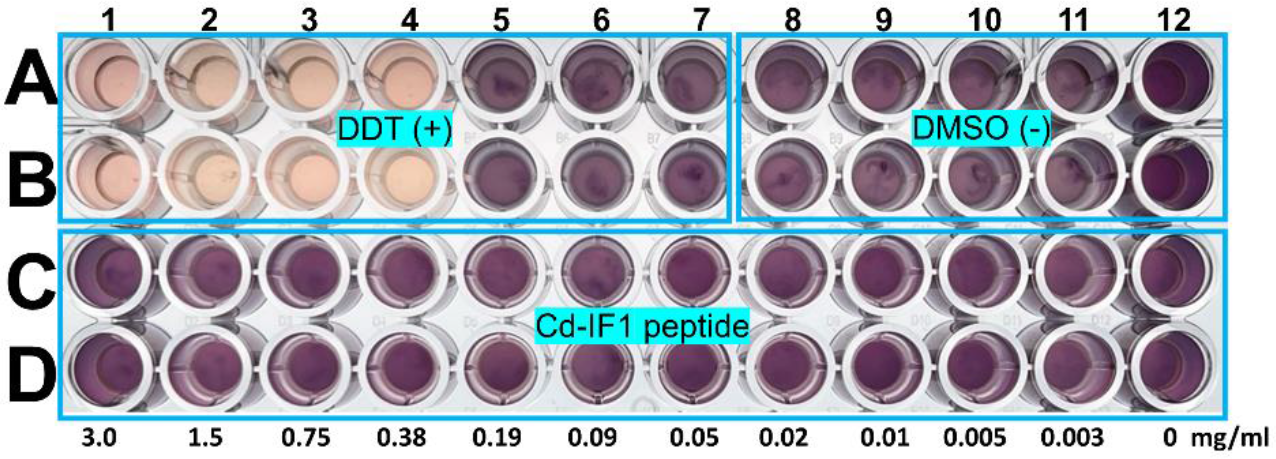
The toxic effect of Cd-IF1 peptide on the growth of human cell cultures tested using HEK-293 cells with positive (+, dichlorodiphenyltrichloroethane (DDT)) and negative (-) control (Dimethyl sulfoxide (DMSO)). DDT at high concentrations (Light pink: A1-A4, B1-B4) inhibits the growth of HEK-293 cells in the Dulbecco’s Modified Eagle Media (DMEM), but not at low concentrations (Dark brown: A5-A7, B5-B7). Cd-IF1 peptide (C1-12, D1-12), like DMSO (A8-A12, B8-B12), does not suppress cell proliferation of HEK-293 in the DMEM at the tested concentrations. The peptide concentrations are labeled under each well.

## Discussion

In this study, we reported the solution structure of *C. difficile* IF1 and its interaction with the 30S ribosomal subunit. *C. difficile* is the most common causative agent of antibiotic-associated diarrhea and gastroenteritis-associated death. The infection caused by *C. difficile* is classified as one of the top five “urgent threats” by the US Centers for Disease Control and Prevention (CDC) [20]. Particularly, CDI has become a growing global concern in the challenging era of COVID-19, urgently requiring the development of efficient treatment and prevention strategies [21]. While numerous studies were on *C. difficile* virulence factors for targeting its toxins [22, 23], this work is focused on protein synthesis in *C. difficile*. There is no structure of *C. difficile* IF1 known to date despite many structures of bacterial IFs being available. The structure of this smallest initiation factor in *C. difficil*e protein synthesis was determined by NMR spectroscopy. Cd-IF1 is comprised of an antiparallel β-barrel of the topology with 5 β-sheets and one short α-helix. The hydrophobic residues, e.g., Valine, Leucine, Isoleucine, and Phenylalanine, are buried in the interior of the barrel to form a hydrophobic core. The polar residues including acidic and basic amino acids, are oriented to the outside of the barrel on the solvent-exposed surface (Figure 7). As a typical OB-fold protein, Cd-IF1 consists of many basic amino acids resulting in its high value of isoelectric point (pI > 9.0). These basic residues are mainly located in Strands 2 and 5 as well as helical region (Figure 7), suggesting this positively charged interface may interact with nucleic acids. However, the posterior on the protein surface is rich in acidic amino acids with negative charges (Figure 2). This asymmetric surface charge distribution likely leading to electric charge attraction and repulsion when close to interacting targets, was thought to be important in stabilizing the binding of IF1 to the ribosome [6].

**Figure 7.**
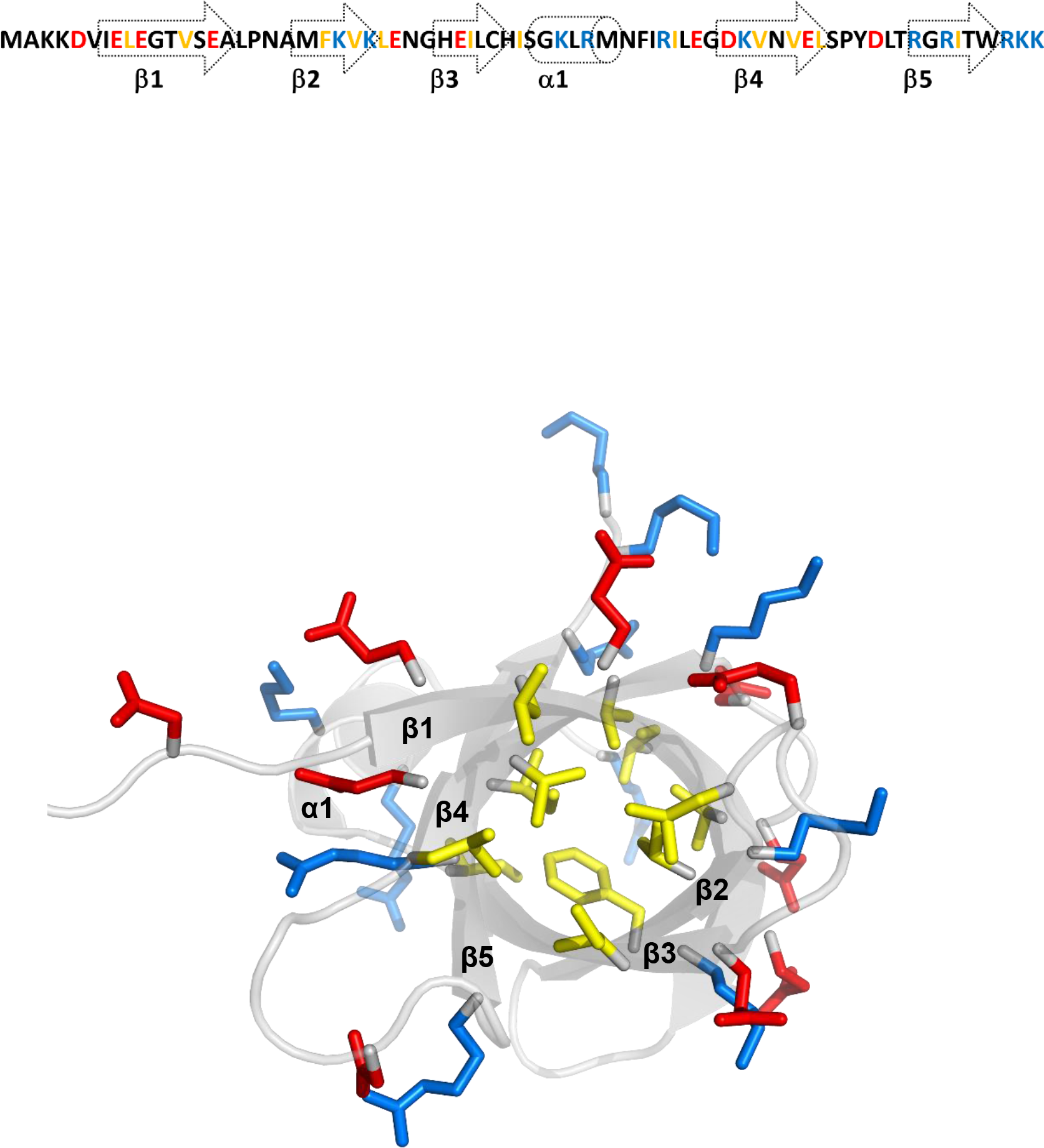
Structural distribution of amino acids in Cd-IF1. Top: the amino acid sequence with secondary structure indicated. Bottom: the NMR structure of Cd-IF1 shows the amino acids buried within the hydrophobic core are highlighted in yellow; the hydrophilic residues on the solvent-exposed surface are in red (acidic) and blue (basic) (bottom view from the long flexible loop harboring the short α-helix)

### Structural comparison of Cd-IF1 with other bacterial IF1 proteins

A number of bacterial IF1 structures have been determined by X-ray crystallography or NMR spectroscopy. As shown in **Figure 8**, these IF1 homologs adopt a compact β-barrel consisting of five β-strands and one short α helix, which is a typical OB fold. The amino acid sequence of Cd-IF1 is most similar to that of *Staphylococcus aureus* (80% identity) but also similar to *Streptococcus pneumoniae* (69%), *Burkholderia thailandensis* (68%), *Mycobacterium tuberculosis* (67%), *Escherichia coli* (65%), *Thermus thermophilus* (65%), and *Pseudomonas aeruginosa* (64%). The main difference in amino acid sequence is observed at both terminal regions-the amino- and carboxyl-terminal ends containing many non-conserved residues. However, the amino acid residues in the long loop region harboring the short α-helix and connecting strands three and four are highly conserved (about 60% sequence identity for all these IF1s). The N-terminal β-strand of these IF1 homologs are very similar as shown in Figure 8. Previous studies showed that *P. aeruginosa* IF1 (PDB 2N78) has an extended β-strand at the C-terminus (β5), which is considerably rigid [9, 24]. The other two IF1s (*S. aureus*, *S. pneumoniae*) share a similar C-terminal structure (PDB 2N8N, 4QL5). The stranded structure of Cd-IF1 C-terminus is shorter than that of *P. aeruginosa* but close to that of *T. thermophilus*. Interestingly, the C-terminal β-strand of *M. tuberculosis* (PDB 3I4O) is the most extended while those of *B. thailandensis* (PDB 2N3S) and *E. coli* (PDB 1AH9) are the shortest, indicative of their high flexibility at the C-terminus. The residues at the C-terminal end (e.g., Arg^70^) have previously been shown critical to IF1 functionality [25]. The structural differences among these IF1 homologs may impact that IF1s from different bacterial species exhibit distinct interaction with their cognate 30S ribosomal subunit.

**Figure 8.**
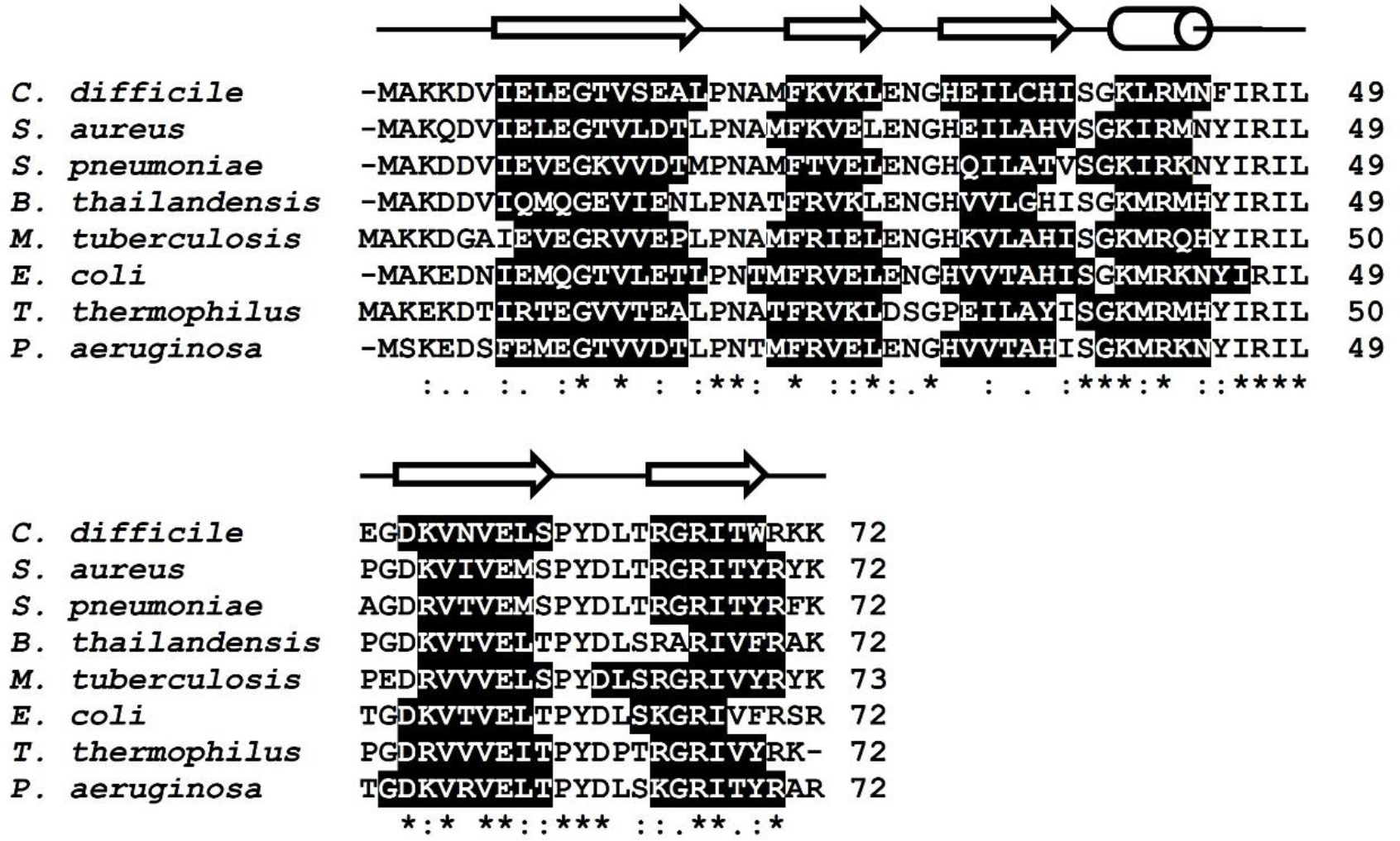
Alignment (Clustal Omega) of the primary sequence of Cd-IF1 (PDB 6C00) with other bacterial homologs from: *S. aureus* (PDB 2N8N), *P. aeruginosa* (PDB 2N78), *E. coli* (PDB 1AH9), *M. tuberculosis* (PDB 3I4O), *S. pneumoniae* (PDB 4QL5), *B. thailandensis* (PDB 2N3S), *T. thermophilus* (PDB 1HR0). Secondary structural elements, highlighted in black, were derived from the PDB structures. The secondary structure elements of Cd-IF1 are indicated schematically above the sequence.

### IF1 functions in bacterial protein biosynthesis

Bacterial IF1 is a highly conserved and indispensable element of the translational apparatus. It plays crucial regulatory roles in translation as well as other cellular processes in prokaryotes. According to the current knowledge, the functions of IF1 include: 1) binds to the 30S ribosomal subunit at the A-site jointly with other components (mRNA, initiator tRNA (fMet-tRNA) and other IFs) to assemble in a 30S initiation complex; 2) modulates the association of IF2 with the ribosome by increasing its binding affinity; 3) prevents the 50S subunit from binding with the 30S and stops the formation of the 70S initiation complex that enters the elongation phase; 4) performs ribosome recycling by working with other initiation factors; 5) binds to RNAs and exerts RNA chaperoning activity [5, 26]. As an OB-fold protein, RNA binding ability is the most important feature of IF1. By binding to the A-site of the 30S subunit, IF1 plays a translation initiation fidelity function by occluding the access of elongator tRNA and other incoming aminoacyl-tRNAs to the A-site [27]. Earlier studies demonstrated IF1, especially in the presence of IF3, inhibits poly-Phe synthesis in the *P. aeruginosa* aminoacylation/translation (A/T) assay due to its ability in preventing association of the ribosomal subunits [9]. These results allowed us to rationally hypothesize that IF1 may inhibit protein synthesis in a high concentration.

Like other bacterial homologs, Cd-IF1 binds with the 30S ribosomal subunit as seen from the NMR titration results (Figure 3). The amino acids with significant perturbations (chemical shift and/or intensity) by the 30S subunits were mainly on one side of the beta-barrel including the short α-helix-involved loop connecting strands three and four, indicative of Cd-IF1 binding interface. The surface of this interface is rich in basic residues (e.g., K39, R41, R46, R64, R66), which can stabilize the binding to the ribosome and is also consistent with earlier structural studies [6]. The structural model of Cd-IF1 in complex with the 30S subunit shows that the short helical region makes the main contact with the groove formed by the ribosomal Helix 44 and Loop 530, which indicates the importance of the helix-harbored long flexible loop between Strand 3 and 4.

### Cd-IF1-derived peptide as a potent antimicrobial agent

The importance of the short α-helix region in Cd-IF1/ribosomal binding by careful inspection of Cd-IF1 interaction with the 30S subunit allowed us to rationally theorize that it may be utilized as an IF1 functional mimic. A Cd-IF1-derived peptide with its amino acid sequence same as the long loop between the third and fourth strand, was designed and synthesized. The peptide was tested to evaluate its inhibitory activity against bacterial growth. As shown in Figure 5, the peptide was able to inhibit the growth of not only *C. difficile* with a MIC of 0.13 mg/ml but also *S. epidermidis* (MIC 0.14 mg/ml) and *E. coli* (MIC 1.12 mg/ml) in the tested media, respectively. In further MIC assays, the peptide demonstrated broad-spectrum antimicrobial activities against both Gram-positive and Gram-negative bacterial strains. Intriguingly, the MIC values for the Gram-positive strains were about 10 times lower than that of the Gram-negative bacteria (Table II), suggesting the peptide is more effective in inhibiting Gram-positive than Gram-negative bacteria. As discussed above, the short α-helix is structurally conserved in bacterial IF1s with a high sequence similarity (Figure 8). It is, therefore, not surprising that the IF1-derived peptide has a wide range of activity against both Gram-positive and Gram-negative organisms.

It should be noted that the peptide was rationally designed as an antimicrobial agent to target bacterial protein synthesis. The structure of the peptide was predicted very similar to the loop between the third and fourth strands in the Cd-IF1 structure (Figure 9). The peptide would bind to the A-site of the 30S subunit like the intact IF1 protein, however, further structural studies are still needed. It should also be noted that this α-helix peptide has five basic residues (Arginine, Lysine, Histidine) in its amino acid sequence (NH_2_-HISGKLRMNFIRILEGDK-COOH). These positive charges may induce bacterial membrane lysis as common antimicrobial peptides do [28]. It is likely that the peptide displays antimicrobial activity via multi-target mode of action-targeting both the ribosomes and membranes.

**Figure 9.**
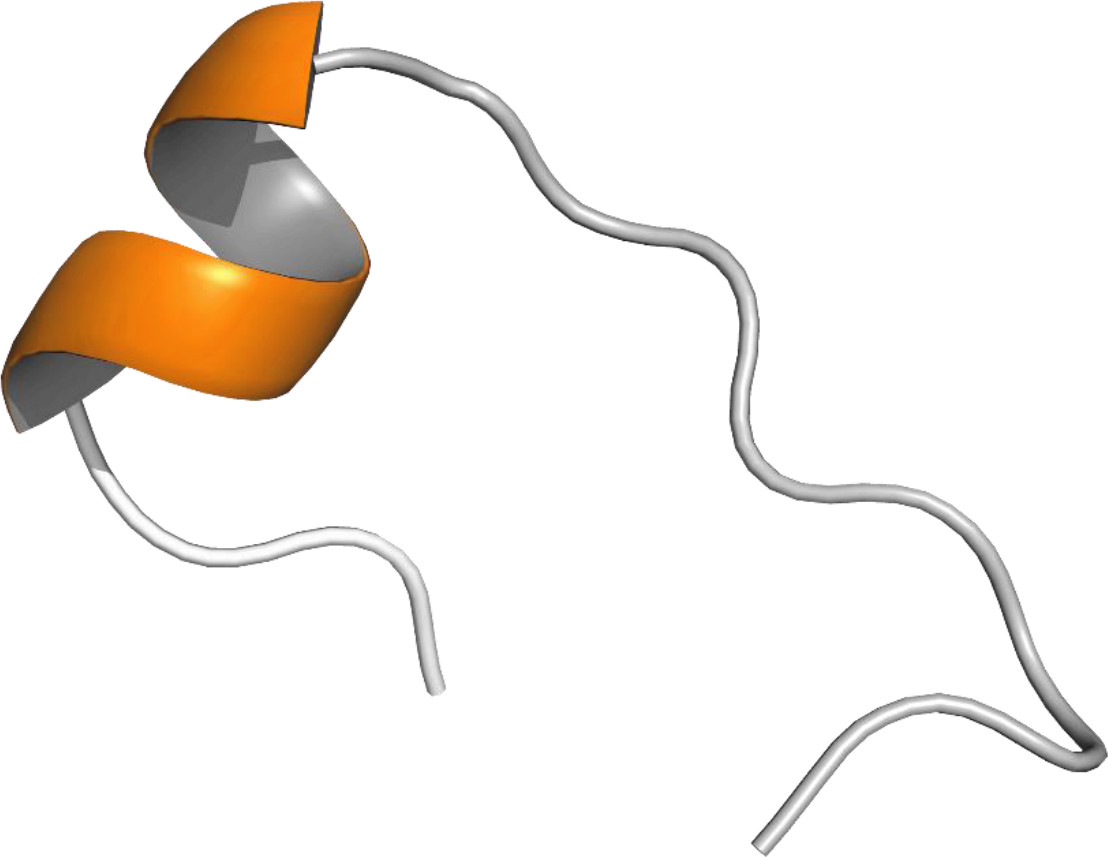
Structure of synthetic Cd-IF1-derived peptide predicted by I-TASSER [32].

An important concern about an antimicrobial agent is cytotoxicity when it is used for antibiotic therapies. Cd-IF1 peptide was tested against the HEK-293 human cell line to see if it could potentially be toxic. After having been treated with the peptides in a concentration up to 3,000 μg/mL for 18 hours, HEK-293 cells were analyzed using MTT assays and showed no observed effect by the peptide compared with control compounds. The peptide did not exhibit toxicity at any concentration tested, suggesting it does not act as an inhibitor like in bacteria to interfere with the growth of eukaryotic cells. Indeed, the peptide was designed based on the structure of IF1 in the bacterium, which is different from its eukaryotic counterpart. Two IF1s (eIF1 and eIF1A) are needed for translation initiation in eukaryotes, despite the binding of eIF1A to the small (40S) ribosomal subunit is near the A-site in a manner similar to the bacterial IF1 [29]. Moreover, the short α-helical region in eIF1A shows low sequence similarity to the Cd-IF1 peptide.

It should be also noted that Cd-IF1 derived peptide exhibits inhibition against the growth of both Gram-positive and Gram-negative bacteria, however, it possesses a relatively moderate antimicrobial activity. A comprehensive structure-activity relationship study is needed to improve its antimicrobial potency. AMPs have recently attracted increasing attention as promising alternative antimicrobial agents because of their unique ability of controlling bacterial infections and low propensity to acquire resistance [3, 28]. Rational design of this peptide may provide a clue for the development of new antimicrobial agents.

## Conclusions

In this study, the structure of translation initiation factor 1 of *C. difficile* was determined by solution NMR methods. The interaction between Cd-IF1 and the 30S ribosomal subunit was studied by NMR titration which allowed the identification of key amino acids involved in the binding with the 30S subunit. The complex structure model of Cd-IF1 and the 30S subunit shows the importance of the short α-helical structure in Cd-IF1 binding to the A-site of 30S subunit. A peptide with the amino acid sequence from the short helical structure of Cd-IF1 was synthesized and tested showing antimicrobial activity. The Cd-IF1 derived peptide inhibits the growth of not only *C. difficile*, but also other Gram-positive and -negative strains. The peptide is likely to be a new generation of antimicrobial peptide candidates.

## Supplementary Materials Author Contributions

Conceptualization, Y.Z.; methodology, E.A., N.B., Y.Z.; software, Y.Z.; validation, Y.Z.; formal analysis, Y.Z.; investigation, E.A., F.A., M.A., Y.Z.; resources, J.B., F.D., Y.Z.; data curation, E.A., F.A., M.A., Y.Z.; writing-original draft preparation, Y.Z.; writing-review and editing, Y.Z.; supervision, Y.Z.; project administration, Y.Z.; funding acquisition, Y.Z. All authors have read and agreed to the published version of the manuscript.

## Institutional Review Board Statement

Not applicable.

## Informed Consent Statement

Not applicable.

## Data Availability Statement

The data that support the findings of this study are available in this manuscript or deposited into public access databases including NMR assignments into the BioMagResBank (http://www.bmrb.wisc.edu/) under Accession Number 27349 NMR structure into the RCSB Protein Data Bank (PDB ID 6C00).

## Acknowledgements

The authors thank Dr. Kristin E. Cano for her technical support and help with NMR experiments at UTHSCSA, and Mr. Thomas Eubanks for his NMR technical support at UTRGV. The authors are grateful to Dr. Daniele Provenzano at UTRGV for valuable suggestions for our IBC protocol approval.

## References

1. Bartlett, J.G., et al., Role of Clostridium difficile in antibiotic-associated pseudomembranous colitis. Gastroenterology, 1978. 75(5): p. 778–82.

2. Blossom, D.B. and L.C. McDonald, The challenges posed by reemerging Clostridium difficile infection. Clin Infect Dis, 2007. 45(2): p. 222–7.

3. Rima, M., et al., Antimicrobial Peptides: A Potent Alternative to Antibiotics. Antibiotics (Basel), 2021. 10(9).

4. Arthithanyaroj, S., et al., Effective inhibition of Clostridioides difficile by the novel peptide CM-A. PLoS One, 2021. 16(9): p. e0257431.

5. Laursen, B.S., et al., Initiation of protein synthesis in bacteria. Microbiol Mol Biol Rev, 2005. 69(1): p. 101–23.

6. Carter, A.P., et al., Crystal structure of an initiation factor bound to the 30S ribosomal subunit. Science, 2001. 291(5503): p. 498-501.

7. Sette, M., et al., The structure of the translational initiation factor IF1 from E.coli contains an oligomer-binding motif. EMBO J, 1997. 16(6): p. 1436–43.

8. Hatzopoulos, G.N. and J. Mueller-Dieckmann, Structure of translation initiation factor 1 from Mycobacterium tuberculosis and inferred binding to the 30S ribosomal subunit. FEBS Lett, 2010. 584(5): p. 1011–5.

9. Hu, Y., et al., Solution structure of protein synthesis initiation factor 1 from Pseudomonas aeruginosa. Protein Sci, 2016. 25(12): p. 2290–2296.

10. Sim, J.H., et al., Optimized Protocol for Simple Extraction of High-Quality Genomic DNA from Clostridium difficile for Whole-Genome Sequencing. J Clin Microbiol, 2015. 53(7): p. 2329–31.

11. Sivashanmugam, A., et al., Practical protocols for production of very high yields of recombinant proteins using Escherichia coli. Protein Sci, 2009. 18(5): p. 936–48.

12. Aguilar, F., N. Banaei, and Y. Zhang, (1)H, (13)C and (15)N resonance assignments and structure prediction of translation initiation factor 1 from Clostridium difficile. Biomol NMR Assign, 2019. 13(1): p. 91-95.

13. Neri, D., et al., Stereospecific nuclear magnetic resonance assignments of the methyl groups of valine and leucine in the DNA-binding domain of the 434 repressor by biosynthetically directed fractional 13C labeling. Biochemistry, 1989. 28(19): p. 7510–6.

14. Delaglio, F., et al., NMRPipe: a multidimensional spectral processing system based on UNIX pipes. J Biomol NMR, 1995. 6(3): p. 277–93.

15. Lee, W., M. Tonelli, and J.L. Markley, NMRFAM-SPARKY: enhanced software for biomolecular NMR spectroscopy. Bioinformatics, 2015. 31(8): p. 1325–7.

16. Nilges, M., et al., Determination of three-dimensional structures of proteins by simulated annealing with interproton distance restraints. Application to crambin, potato carboxypeptidase inhibitor and barley serine proteinase inhibitor 2. Protein Eng, 1988. 2(1): p. 27-38.

17. Schrodinger, L., The PyMOL Molecular Graphics System. 2010.

18. Murzin, A.G., OB(oligonucleotide/oligosaccharide binding)-fold: common structural and functional solution for non-homologous sequences. EMBO J, 1993. 12(3): p. 861–7.

19. Valdez, N., et al., Rational Design of an Antimicrobial Peptide Based on Structural Insight into the Interaction of Pseudomonas aeruginosa Initiation Factor 1 with Its Cognate 30S Ribosomal Subunit. ACS Infect Dis, 2021. 7(12): p. 3161–3167.

20. Control, C.f.D. and Prevention, Antibiotic resistance threats in the United States, 2019. 2019: US Department of Health and Human Services, Centres for Disease Control and….

21. Spigaglia, P., Clostridioides difficile infection (CDI) during the COVID-19 pandemic. Anaerobe, 2022. 74: p. 102518.

22. Awad, M.M., et al., Clostridium difficile virulence factors: Insights into an anaerobic spore-forming pathogen. Gut Microbes, 2014. 5(5): p. 579–93.

23. Chen, P., et al., Structural basis for CSPG4 as a receptor for TcdB and a therapeutic target in Clostridioides difficile infection. Nature Communications, 2021. 12(1): p. 3748.

24. Bernal, A., et al., (1)H, (13)C and (15)N resonance assignments and secondary structure analysis of translation initiation factor 1 from Pseudomonas aeruginosa. Biomol NMR Assign, 2016. 10(2): p. 249-52.

25. Spurio, R., et al., Site-directed mutagenesis and NMR spectroscopic approaches to the elucidation of the structure-function relationships in translation initiation factors IF1 and IF3. Biochimie, 1991. 73(7-8): p. 1001–6.

26. Croitoru, V., et al., RNA chaperone activity of translation initiation factor IF1. Biochimie, 2006. 88(12): p. 1875–1882.

27. Moazed, D., et al., Specific protection of 16 S rRNA by translational initiation factors. J Mol Biol, 1995. 248(2): p. 207–10.

28. Lei, J., et al., The antimicrobial peptides and their potential clinical applications. Am J Transl Res, 2019. 11(7): p. 3919–3931.

29. Lomakin, I.B. and T.A. Steitz, The initiation of mammalian protein synthesis and mRNA scanning mechanism. Nature, 2013. 500(7462): p. 307-311.

30. Jurrus, E., et al., Improvements to the APBS biomolecular solvation software suite. Protein Sci, 2018. 27(1): p. 112–128.

31. Zhang, Y., et al., Backbone chemical shift assignments of mouse HOXA13 DNA binding domain bound to duplex DNA. Biomol NMR Assign, 2010. 4(1): p. 97–9.

32. Yang, J. and Y. Zhang, I-TASSER server: new development for protein structure and function predictions. Nucleic Acids Res, 2015. 43(W1): p. W174–81.

